# Investigation of risk factors associated with *Ancylostoma* spp. infection and the benzimidazole F167Y resistance marker polymorphism in dogs from the United States

**DOI:** 10.1101/2024.09.08.611871

**Authors:** Pablo D. Jimenez Castro, Jennifer L. Willcox, Haresh Rochani, Holly L. Richmond, Heather E. Martinez, Cecilia E. Lozoya, Christian Savard, Christian M. Leutenegger

## Abstract

*Ancylostoma caninum* is the most significant intestinal nematode parasite of dogs. We acquired fecal surveillance data using molecular diagnostics in a large population of dogs in the United States (US). A diagnostic test using real-time PCR (qPCR) for *Ancylostoma* spp. and allele-specific qPCR detecting the SNP F167Y was used in 885,424 canine fecal samples collected between March 2022 and December 2023. Overall, *Ancylostoma* spp. had a prevalence of 1.76% (15,537/885,424), with the highest observed in the South 3.73% (10,747/287,576), and the lowest in the West 0.45% (632/140,282). Within the subset of *Ancylostoma* spp*.-*detected dogs used for further analysis, the F167Y SNP had an overall prevalence of 14.2% with the highest in the West and the lowest in the Midwest (10.76%). The greyhound breed exhibited a higher prevalence of *Ancylostoma* spp. infections (17.03%) and a higher prevalence of the F167Y polymorphism (33.6%) compared to non-greyhound breeds (13.7% and 2.08%), respectively, but were not associated with the highest breed risk for the F167Y polymorphism. Sex did not influence hookworm infection nor F167Y polymorphism prevalence. Intact dogs had a prevalence of hookworm infection and F167Y polymorphism of 2.51% and 14.6%, respectively. Puppies showed increased prevalence of hookworms (3.70%) and the F167Y SNP (17.1%). Greyhounds, bluetick coonhounds, and boerboels had the highest relative risks (RR) for hookworm infection, while Cavalier King Charles spaniels, Havanese, and shiba inus had the lowest. The top and bottom three with the highest and lowest RR for the F167Y SNP were the old English sheepdog, American foxhound, and toy poodle Toy, and shih tzu, Maltese, and Australian cattle dogs, respectively. This study highlights the value of an accessible diagnostic qPCR test with fast turnaround times in unraveling the molecular epidemiology of hookworms and benzimidazole resistance, as well as explore potentially important risk factors associated with infection in medicalized dogs.

**Highlights:** - Greyhounds had the highest RR relative risk for *Ancylostoma* spp., but only fourth for the *A. caninum* F167Y polymorphism.

- The highest prevalence of *Ancylostoma* spp. in the United States was in the South.

- The West had the lowest prevalence for *Ancylostoma* spp., but the highest prevalence for the *A. caninum* F167Y SNP.

- Puppies had the highest prevalence and AOR for *Ancylostoma* spp. and F167Y polymorphism.

## 1. Introduction

The canine hookworm, *Ancylostoma caninum*, is the most significant intestinal nematode parasite of dogs in the United States (US), with the prevalence dependent on factors such as age, level of care, and geographic location (Little et al., 2009). Concurrently, there has been a steady increase in reported canine hookworm frequencies from 2012 to 2018, indicating an overall increase of 45.6% over the seven-year period (Drake and Carey, 2019). Furthermore, along with *Giardia duodenalis*, hookworms had the highest positive proportion in canine fecal samples submitted to a veterinary reference laboratory from 2017 to 2019 (Sweet et al., 2021). In a recent retrospective study assessing results from fecal flotation of dogs across 10 veterinary diagnostic laboratories in the US, Ancylostomatidae eggs exhibited a prevalence of 5.63% (Sobotyk et al., 2021). The risk for widespread anthelmintic drug resistance in companion animals, in contrast to livestock and horses, has surfaced only recently, with resistance found against the three primary drug classes used for treating parasitic nematodes of dogs in the US (Jimenez Castro et al., 2019; Von Samson-Himmelstjerna et al., 2021). In most of these cases, the F167Y (TTC > TAC) single nucleotide polymorphism (SNP) in the *A. caninum* β-tubulin isotype 1 gene polymorphism was associated with treatment resistance to benzimidazole anthelmintics (Kitchen et al., 2019; Jimenez Castro et al., 2019; Venkatesan et al., 2023). Subsequently, several cases have revealed the presence of benzimidazole treatment-resistant hookworms, particularly in greyhounds, which also harbored multiple anthelmintic drug resistant (MADR) *A. caninum* infections (Jimenez Castro et al., 2020; Jimenez Castro et al., 2021; Jimenez Castro et al., 2022). Awareness of treatment-resistant hookworms has not been limited to the US, with reports dating back to 1987 indicating pyrantel treatment failure against *A. caninum* in Australia, along with subsequent reports of SNPs associated with benzimidazole resistance in Brazil (Jackson et al., 1987; Hopkins et al., 1988; Hopkins and Gyr, 1991; Kopp et al., 2007; Furtado et al., 2014; Furtado and Rabelo, 2015). More recent reports have further substantiated the prevalence of resistant *A. caninum* infections across various non-greyhound dog breeds (Jimenez Castro et al., 2022; Venkatesan et al., 2023; Leutenegger et al., 2023a,). Taken together, these observations indicate a broader than anticipated spread of resistant hookworm infections, emphasizing the urgency for readily accessible and cost-effective drug resistance screening tests for veterinary wellness programs (Marsh and Lakritz, 2023). Interestingly, a recent study delving deeper into the data generated by Venkatesan et al., 2023, indicated that *A. braziliense, A. tubaeforme*, and *Uncinaria stenocephala* sequences did not include the benzimidazole resistance-associated SNPs (Stocker et al., 2024).

Given the confirmed presence of the F167Y polymorphism, among greyhound populations, alongside recent reports of resistant isolates in the broader pet dog population (Jimenez Castro et al, 2022; Venkatesan et al., 2023; Leutenegger et al., 2023a), it becomes imperative to devise control strategies and treatment guidelines. These initiatives are crucial for upholding the efficacy of current drugs and monitoring the emergence of resistance to additional anthelmintic compounds and will require the use of diagnostic protocols for widespread screening in pet wellness settings.

Notably, we recently validated a hydrolysis probe-adapted allele-specific qPCR test, ensuring high analytical specificity for the F167Y SNP (TTC > TAC) (Leutenegger et al., 2023a). This test showed a sensitivity of 4 molecules per reaction, surpassing the commonly required sensitivity of 10 molecules per reaction. Similarly, the *Ancylostoma* spp.-specific qPCR exhibited high analytical sensitivity, resulting in a limit of detection of 6.33 eggs per gram of feces (Leutenegger et al., 2023a). In a recent study, using greater than 900 fecal samples from dogs and cats submitted to a veterinary reference laboratory for centrifugal floatation with zinc sulfate (specific gravity of 1.18 ± 0.005), the samples were then evaluated by a broad qPCR panel (KeyScreen^®^ GI Parasite PCR, Antech Diagnostics, Fountain Valley, CA). A significantly higher parasite frequency and almost three times the co-infections were detected by the qPCR panel (Leutenegger et al., 2023b). Moreover, the qPCR had an increased ability to detect the parasites where the centrifugal floatation had a reduced or had no detection, namely *Giardia duodenalis*, *Trichuris* spp., *Cystoisospora* spp., *Tritrichomonas blagburni*, and *Toxoplasma gondii* (Leutenegger et al., 2023b).

The American Association of Veterinary Parasitologists (AAVP) recommends using the term “MADR” to specifically describe infections that persist despite treatment with two or more commonly used anthelmintic drugs for patent *A. caninum* infections following an appropriate diagnostic workup (i.e., sample collected the day of treatment and 14 days later). It is important to avoid using the acronym “MDR” for multiple drug resistance in this context, as it could be confused with the terminology related to the MDR1 gene.

To date, there have been few studies in North America assessing the prevalence of the *A. caninum* F167Y SNP by geographical location, breed, age, and even seasonality (Venkatesan et al., 2023 Leutenegger et al., 2024). Furthermore, there is a knowledge gap in terms of the actual relative risk and odds ratio of the resistant infections to which there are molecular markers.

The primary objective of our fecal surveillance study was to offer insights into the molecular epidemiology of hookworm infection and the *A. caninum* F167Y benzimidazole resistant polymorphism. Specifically, to evaluate the prevalence and relative risk association for patient characteristics (age, sex, neuter status, breed, and geographic location) using a validated and published molecular diagnostic qPCR test (Leutenegger et al., 2023b).

## 2. Materials and methods

### 2.1 Data collection

A subset of data of 885,424 canine fecal samples were collected between March 2022 and December 2023, from the laboratory information management system of a commercial reference laboratory (Antech Diagnostics, Inc.) as part of a larger parasite molecular diagnostic PCR panel (KeyScreen^®^ GI Parasite PCR, Antech Diagnostics, Fountain Valley, CA) (Leutenegger et al., 2023b). Part of this data has already been descriptively reported (Leutenegger et al., 2024). However, in the previous study we did not perform any statistical analyses evaluating the relative risks nor the adjusted odds ratio for the different patient characteristics. Canine ages were divided into four groups: puppies aged less than a year, young adults aged 1-3 years, mature adults aged 4-7 years, and senior dogs aged over 7 years as per recent American Animal Hospital Association guidelines (Creevy et al., 2019). In cases where breed information was added as free text by the veterinarian or a member of the veterinary staff, breed information was collected as per the entry into the clinic software. Geographically, the data was divided as per the US Census Bureau into 4 regions: the West, Midwest, South and Northeast, as is commonly described (https://data.census.gov/). This geographical division has been previously used in other prevalence survey studies (Little et al., 2009, Venkatesan et al., 2023, Leutenegger et al., 2024).

### 2.2. Real-time PCR tests and quality controls

The data set for this study was obtained with a commercially available GI parasite molecular test, which identifies 20 individual parasites, the F167Y polymorphism of *A. caninum* associated with resistance to benzimidazoles, and *Giardia duodenalis* with zoonotic potential (KeyScreen^®^ GI Parasite PCR, Antech Diagnostics, Fountain Valley, CA, (Leutenegger et al., 2023b)). Even though the assay does not differentiate between the different *Ancylostoma* species that could infect dogs in the US, recent widespread surveys performed in the US have shown that the most common species found is *Ancyslostoma caninum* (Venkatesan et al., 2023, Stocker et al., 2024). Additionally, two quality controls were used in conjunction with every diagnostic sample, as previously described (Leutenegger et al., 2024).

### 2.3. Statistical Methods

Descriptive statistics were employed to summarize the characteristics of the study population, including demographic information and prevalence rates of *Ancylostoma* spp. and *A. caninum* F167Y infections. Measures such as frequencies and percentages were utilized to describe the variability of the data. Relative risks (RRs) were calculated to assess the association between breed as a risk factor and *Ancylostoma* spp*. (*or with *A. caninum* F167Y) infection. The RRs were estimated along with their corresponding 95% confidence intervals (CIs) using standard methods. Relative risk and odds ratios in which the 95% CI did not include 1 were considered statistically significant.

Logistic regression analysis was performed to further investigate the relationship between predictor variables and the likelihood of *Ancylostoma* spp. (or with *A. caninum* F167Y) infection while adjusting for potential confounding factors. The outcome variable was the presence or absence of *Ancylostoma* spp*. (*or with *A. caninum* F167Y) infection, and predictor variables included neuter status (yes/no), sex (male/female), region of residence (categorical), and age (categorical). Adjusted odds ratios (AORs) along with their 95% CIs were calculated to determine the strength and direction of association between each predictor variable and the outcome variable, and these were considered statistically significant if these did not include 1. We chose not to explore all possible interactions, mainly to maintain model stability, and because the interaction between region and age seemed more meaningful to analyze. All statistical analyses were performed using SAS version 9.4.

## 3. Results

### 3.1 Regional distribution of Ancylostoma spp. and A. caninum F167Y

The study investigated the prevalence of *Ancylostoma* spp. infection, particularly focusing on the *A. caninum* F167Y mutation, across different demographic categories in a large dataset of veterinary submissions. The results, presented in Table 1, provide valuable insights into the epidemiology of this parasite, as well as subpopulations with the presence of the benzimidazole resistance associated F167Y polymorphism.

**Table 1:** Number of total submissions for *Ancylostoma* spp. and the *A. caninum* F167Y Polymorphism Compared to the Total or Within Sample Set Across Various Demographic Categories.

|  | *Total Submissions* |  | *Ancylostoma* spp. |  |  | *A. caninum* F167Y |  |  |
| --- | --- | --- | --- | --- | --- | --- | --- | --- |
|  | n= | % | n= | %<sup>a</sup> | %<sup>b</sup> | n= | %<sup>a</sup> | %<sup>c</sup> |
| **Age Category** |  |  |  |  |  |  |  |  |
| Puppy (<1y) | 214,146 | 30.45% | 7,940 | 51.92% | 3.71% | 1,360 | 62.41% | 17.13% |
| Young adult (1-3y) | 126,462 | 17.98% | 3,014 | 19.71% | 2.38% | 378 | 17.35% | 12.54% |
| Mature adult (4-7y) | 173,752 | 24.71% | 2,793 | 18.26% | 1.61% | 325 | 14.92% | 11.64% |
| Senior (=>7y) | 188,894 | 26.86% | 1,544 | 10.10% | 0.82% | 116 | 5.32% | 7.51% |
| **Sex** |  |  |  |  |  |  |  |  |
| Female | 353,101 | 48.63% | 7,437 | 48.21% | 2.11% | 1,044 | 47.76% | 14.04% |
| Male | 368,285 | 50.72% | 7,837 | 50.79% | 2.13% | 1,119 | 51.19% | 14.28% |
| Unknown | 4,735 | 0.65% | 154 | 1.00% | 3.25% | 23 | 1.05% | 14.94% |
| **Neuter Status** |  |  |  |  |  |  |  |  |
| No | 232,831 | 31.70% | 9,035 | 57.58% | 3.88% | 1,415 | 63.82% | 15.66% |
| Yes | 501,543 | 68.30% | 6,656 | 42.42% | 1.33% | 802 | 36.18% | 12.05% |
| **US Region** |  |  |  |  |  |  |  |  |
| Northeast | 162,606 | 23.84 % | 2,399 | 15.79% | 1.48% | 358 | 16.58% | 14.92% |
| Midwest | 91,634 | 13.43 % | 1,412 | 9.29% | 1.54% | 152 | 7.04% | 10.76% |
| South | 287,576 | 42.16 % | 10,747 | 70.75% | 3.74% | 1,524 | 70.59% | 14.18% |
| West | 140,282 | 20.57 % | 632 | 4.16% | 0.45% | 125 | 5.79% | 19.78% |
| **Predispose Status** |  |  |  |  |  |  |  |  |
| Greyhound | 1,902 | 0.26% | 324 | 2.06% | 17.03% | 109 | 4.92% | 33.64% |
| Non-Greyhound | 737,207 | 99.74% | 15,367 | 97.94% | 2.08% | 2,108 | 95.08% | 13.72% |

<sup>a</sup> % of cases relative to the total number of submissions for each respective category.
<sup>b</sup> % of the specific age category, sex, neuter status, region, or predispose status that is infected with *Ancylostoma* spp. out of the total number of submissions in that category.
<sup>c</sup> % of the specific age category, sex, neuter status, region, or predispose status that is infected with *A. caninum* F167Y out of the total number of infections with *Ancylostoma* spp. within that category.

Across the study population, the prevalence was 1.765% (15,537/885,424) for *Ancylostoma* spp. and 14.442% (2,244/15,537) for the *A. caninum* F167Y polymorphism. Of these, 15,190 samples contained adequate geographic and demographic information and were used for further analysis and mapping. Geographic variations in prevalence were observed across different regions of the US. The South exhibited the highest overall prevalence of *Ancylostoma* spp. infection and third in *A. caninum* F167Y detected proportion, with 3.74% (10,747/287,576) and 14.18% (1524/10,747), respectively. Conversely, the West region had the lowest prevalence rate for *Ancylostoma* spp. infection (0.45% (632/140,282) but the highest prevalence for *A. caninum* F167Y with 19.78% (125/632), suggesting regional disparities in susceptible and benzimidazole-resistant hookworm distribution.

### 3.2 Sex and Age distribution of Ancylostoma spp. and A. caninum F167Y

In terms of age distribution, puppies (<1 year) exhibited the highest proportion of submissions. Within this age category, *Ancylostoma* spp. infection in puppies (<1 year) represented the group both with the highest proportion of infections (51.92%) and prevalence (3.71% (7,940/214,146)). Puppies also had the highest proportion of *A. caninum* F167Y infections with 17.13% (1,360/7,940). Notably, the prevalence inversely decreased with increasing age, with senior dogs (≥7 years) showing the lowest at 0.82% (1,544/188,894), even though this age group represented 26.86% of submissions. However, even within this age group, *A. caninum* F167Y remained detectable at 7.51% (116/1,544). Regarding sex, a balanced distribution was observed, with females comprising 48.63%, and males 50.72% of total submissions. Similar prevalence rates of *Ancylostoma* spp. were found among both sexes, with 2.11% (7,437/353,101) in females and representing 48.21% of submissions, and 2.13% (7,837/368,285) in males and representing 50.79% of submissions.

### 3.3 Breed and neuter status distribution of Ancylostoma spp. and A. caninum F167Y

Analysis by neuter status revealed that neutered animals accounted for the majority of submissions (68.30% (501,543/734,374)) and displayed a lower prevalence of *Ancylostoma spp.* compared to unneutered animals (1.33% (6,656/501,543) vs 3.88% (9,035/232,831)). This finding suggests a potential association between reproductive status and higher risk of exposure or susceptibility to *Ancylostoma spp.* infection. Additionally, unneutered animals exhibited a higher prevalence of *A. caninum* infection (15.66% (1,415/9,035)) compared to neutered animals (12.05% (802/6,656)), further supporting the association between reproductive status and the risk of parasitic infections. Lastly, predisposing factors such as breed were examined, with greyhounds accounting for only 0.26% of the total submissions that included breed information but showing a notably higher prevalence for both *Ancylostoma* spp. and *A. caninum* F167Y infections compared to non-greyhound breeds (17.03% (324/1,902) vs. 2.08% (15,367/737,207)) and (33.64% (109/324) vs. 13.71% (2108/15,367)), respectively.

### 3.4 Relative risks analysis of Ancylostoma spp. and A. caninum F167Y infection

The results of the study, presented in Table 2, demonstrate the relative risk (RR) of *Ancylostoma* spp. infection and presence of the *A. caninum* F167Y polymorphism from the top 20 breeds, along with their corresponding 95% confidence intervals (CI).

**Table 2:** The Relative Risk (RR) of *Ancylostoma* spp. and *A. caninum* F167Y for the top 20 breeds.

| *Ancylostoma* spp. |  |  | *A. caninum* F167Y |  |  |
| --- | --- | --- | --- | --- | --- |
| Breed | RR | 95% CI | Breed | RR | 95% CI |
| Greyhound | 8.03 | (7.27, 8.88) | Old English sheepdog | 3.79 | (2.36, 6.09) |
| Bluetick coonhound | 6.79 | (4.96, 9.29) | American foxhound | 3.55 | (2.23, 5.65) |
| Boerboel | 5.86 | (2.94, 11.68) | Poodle, toy | 3.20 | (1.97, 5.20) |
| Field spaniel | 4.40 | (2.37, 8.18) | Greyhound | 2.45 | (2.10, 2.87) |
| Bloodhound | 4.07 | (2.96, 5.61) | Bluetick coonhound | 2.15 | (1.28, 3.61) |
| American foxhound | 3.70 | (2.38, 5.76) | Bernese mountain dog | 2.00 | (1.31, 3.03) |
| Treeing walker coonhound | 3.49 | (2.24, 5.43) | Shetland sheepdog | 1.99 | (1.06, 3.73) |
| Great Pyrenees | 3.17 | (2.76, 3.64) | English springer spaniel | 1.96 | (1.08, 3.53) |
| American black & tan coonhound | 2.92 | (2.31, 3.70) | Poodle | 1.95 | (1.59, 2.38) |
| Cane Corso | 2.59 | (2.10, 3.19) | Beagle | 1.60 | (1.31, 1.95) |
| American pitbull terrier | 2.55 | (2.09, 3.12) | Jack Russell terrier | 1.60 | (1.06, 2.41) |
| American Staffordshire terrier | 2.10 | (1.84, 2.39) | Cane Corso | 1.56 | (1.03, 2.35) |
| Staffordshire bull terrier | 2.10 | (1.74, 2.52) | American black & tan coonhound | 1.53 | (0.96, 2.44) |
| Catahoula leopard dog | 2.07 | (1.63, 2.63) | Saint Bernard | 1.52 | (0.75, 3.09) |
| Boykin spaniel | 2.04 | (1.20, 3.48) | Bull terrier | 1.47 | (0.72, 2.99) |
| Redbone coonhound | 1.91 | (0.97, 3.77) | Bloodhound | 1.46 | (0.75, 2.83) |
| Bulldog | 1.88 | (1.63, 2.16) | Cocker spaniel | 1.42 | (0.85, 2.36) |
| Affenpinscher | 1.85 | (1.29, 2.66) | Australian shepherd, miniature | 1.35 | (0.72, 2.52) |
| Great Dane | 1.83 | (1.56, 2.15) | Bulldog | 1.32 | (0.98, 1.79) |
| Doberman pinscher | 1.83 | (1.55, 2.15) | Pembroke Welsh corgi | 1.31 | (0.75, 2.30) |

Of the breeds with the most elevated RRs, greyhounds exhibited the highest RR for *Ancylostoma* spp. infection (RR = 8.03, 95% CI: 7.27-8.88) and fourth highest for the *A. caninum* F167Y polymorphism (RR = 2.45, 95% CI: 2.10-2.87). This suggests a significantly elevated risk or history of exposure in this breed compared to others. Similarly, breeds such as bluetick coonhound, boerboel, and field spaniel also displayed significantly higher RRs for *Ancylostoma* spp. infection, indicating an increased risk of exposure within these populations. Among the breeds examined, the old English sheepdog demonstrated the highest RR for *A. caninum* F167Y (RR = 3.79, 95% CI: 2.36-6.09), followed by the American foxhound (RR = 3.55, 95% CI: 2.23-5.65) and the toy poodle (RR = 3.20, 95% CI: 1.97- 5.20). Conversely, breeds such as the beagle, jack russell terrier, and cocker spaniel exhibited lower RRs for both *Ancylostoma* spp. infection and the *A. caninum* F167Y polymorphism, indicating a comparatively reduced risk within these populations. Figure 2 and Figure 3 illustrate the results from Table 2 graphically, providing a visual comparison of the RR results. These findings highlight breed-specific differences in risk of exposure to the *A. caninum* F167Y polymorphism, suggesting potential genetic or physiological factors at play.

**Figure 1.**
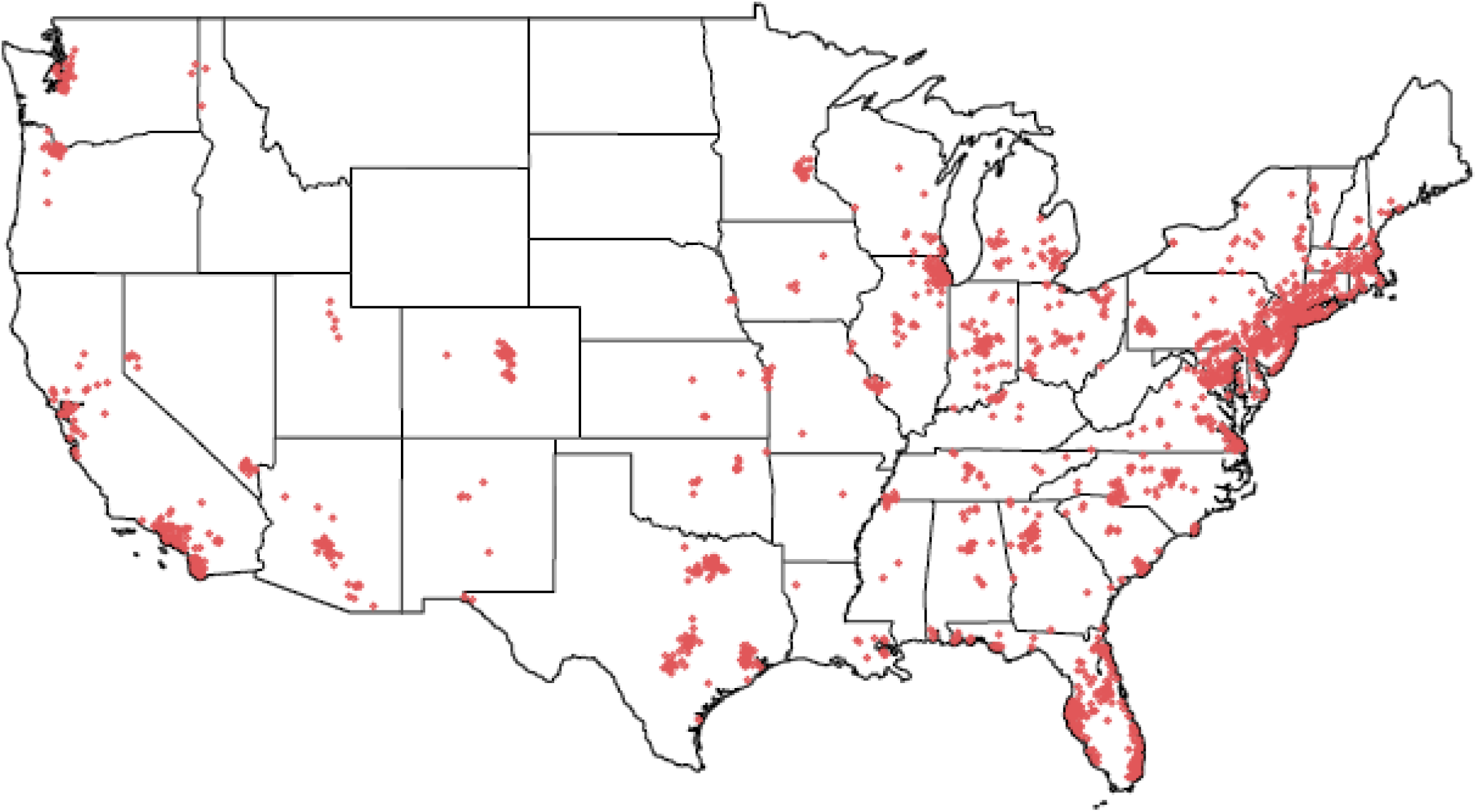
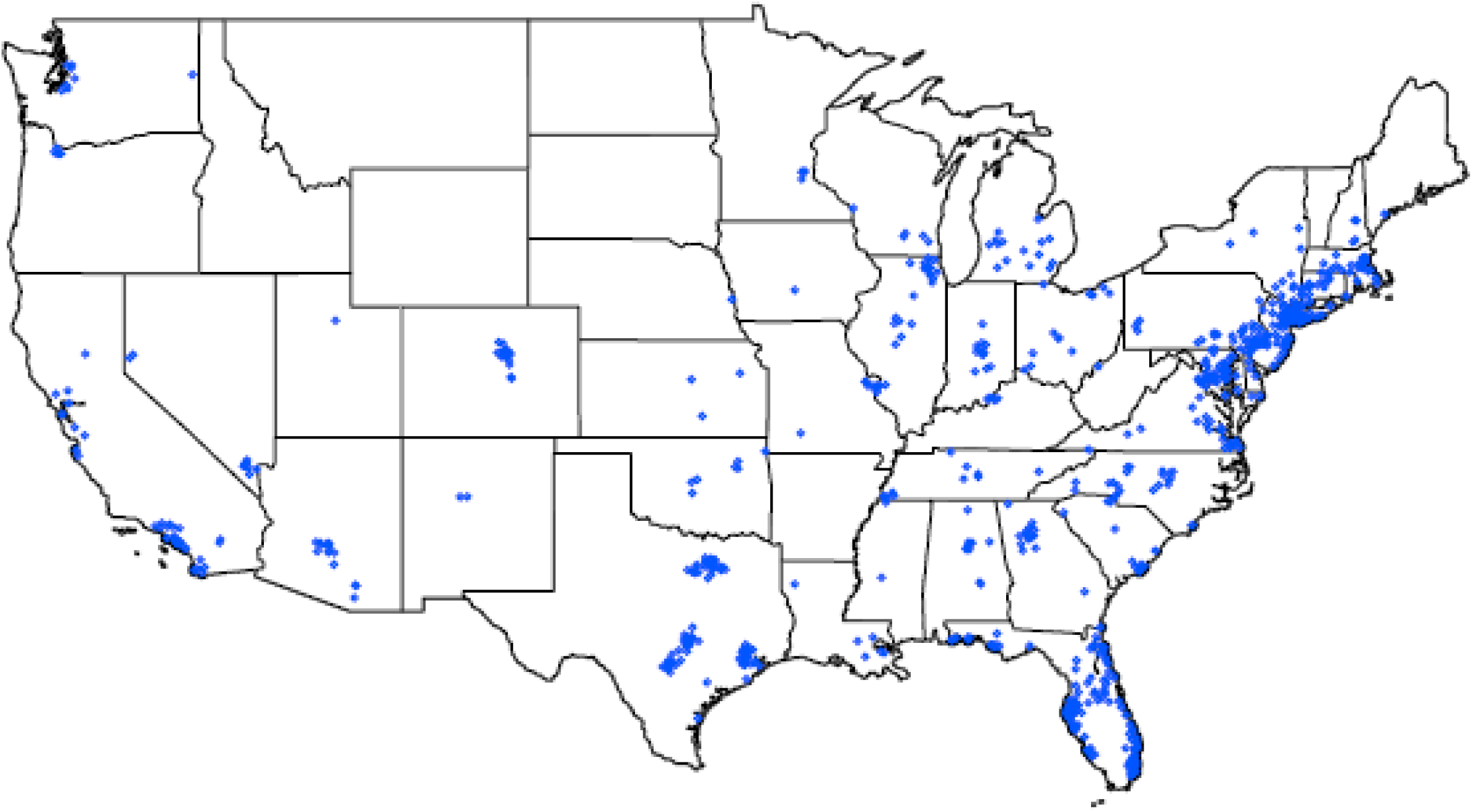
A: Distribution of *Ancylostoma spp.* in the US. B: Distribution of benzimidazole resistance associated F167Y polymorphism in *A. caninum* in the US with this cohort.

**Figure 2.**
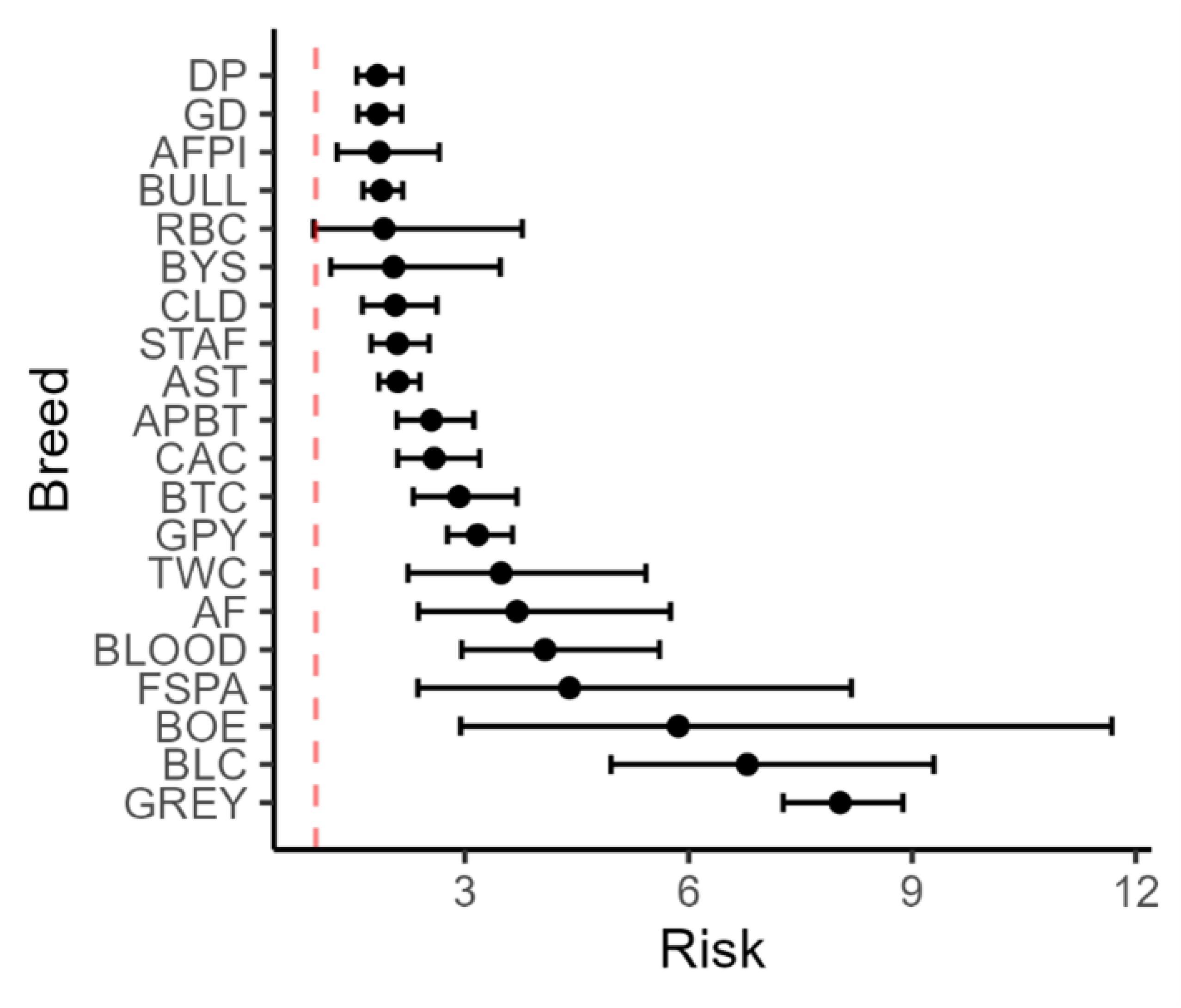
Relative risks by Top 20 Dog Breed for *Ancylostoma* spp.

**Figure 3.**
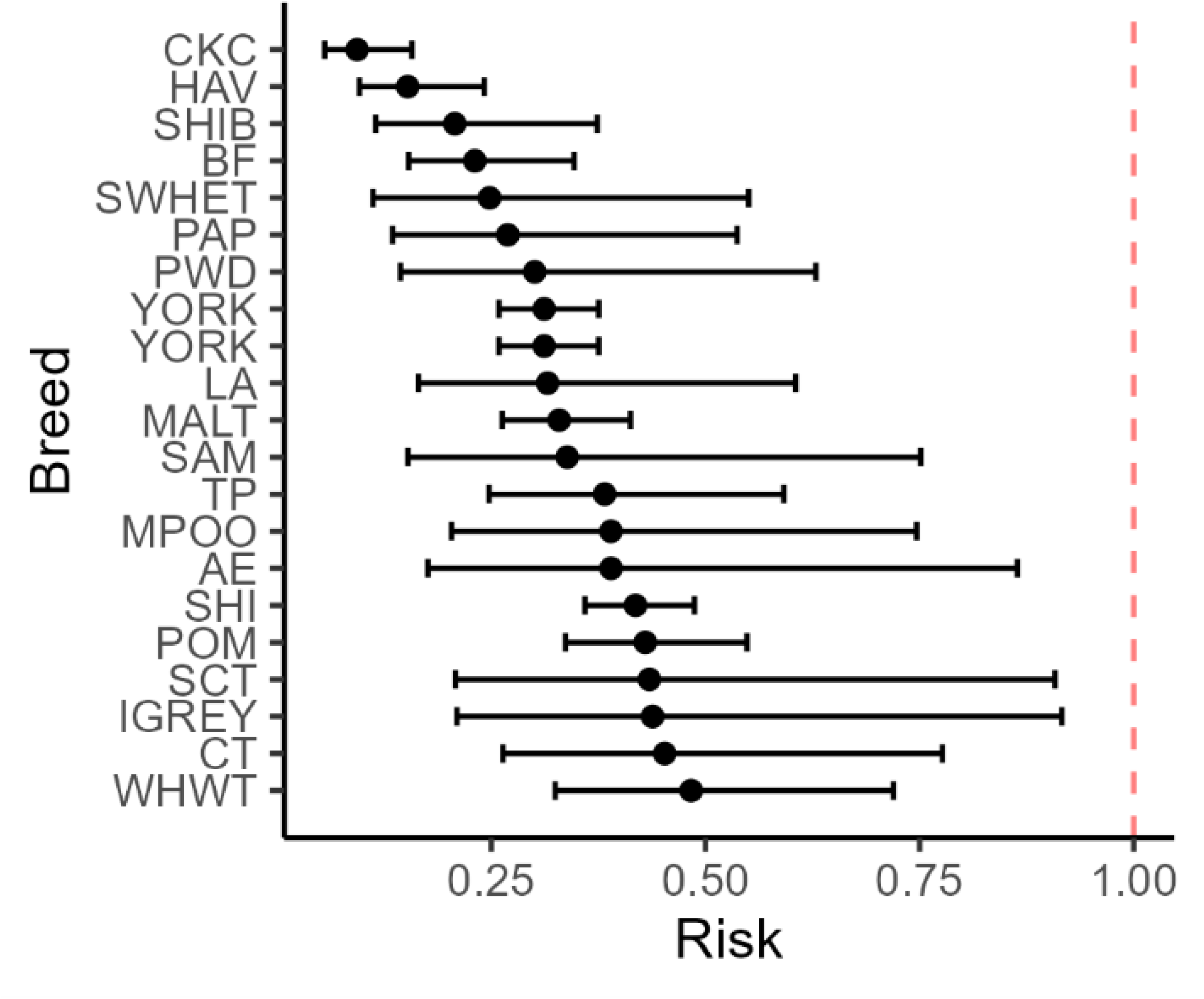
Relative risks by Top 20 Dog Breed for *A. caninum* F167Y.

Table 3 presents the RR of *Ancylostoma* spp. infection and the *A. caninum* F167Y polymorphism for the bottom 20 dog breeds, along with their corresponding 95% CI. Among the breeds with the lowest RR for *Ancylostoma* spp. infection, the cavalier King Charles spaniel exhibited the lowest risk (RR = 0.09, 95% CI: 0.06-0.16), followed by the Havanese (RR = 0.15, 95% CI: 0.10-0.24) and the shiba inu (RR = 0.21, 95% CI: 0.11-0.37). For the *A. caninum* F167Y polymorphism, the Great Pyrenees displayed the lowest RR (RR = 0.40, 95% CI: 0.23-0.72), indicating a reduced risk compared to other breeds. Other breeds such as the Staffordshire bull terrier (RR = 0.78, 95% CI: 0.46-1.33) and the Belgian Malinois (RR = 0.79, 95% CI: 0.39-1.58) demonstrated slightly higher RRs for *Ancylostoma* spp. infection, although still within a relatively low-risk range (RRs are still < 1). Some breeds, such as the American Staffordshire terrier exhibited an RR close to 1 (RR = 1.01, 95% CI: 0.73-1.40), suggesting a comparable risk to the overall population. For the *A. caninum* F167Y polymorphism, breeds such as the Chihuahua (RR = 0.83, 95% CI: 0.60-1.15) and the Labrador retriever (RR = 0.85, 95% CI: 0.71-1.02) demonstrated RRs close to 1, indicating as well a similar risk profile to the overall population. Figure 4 and Figure 5 illustrate the results from Table 3 graphically, providing a visual comparison of the RR results.

**Figure 4:**
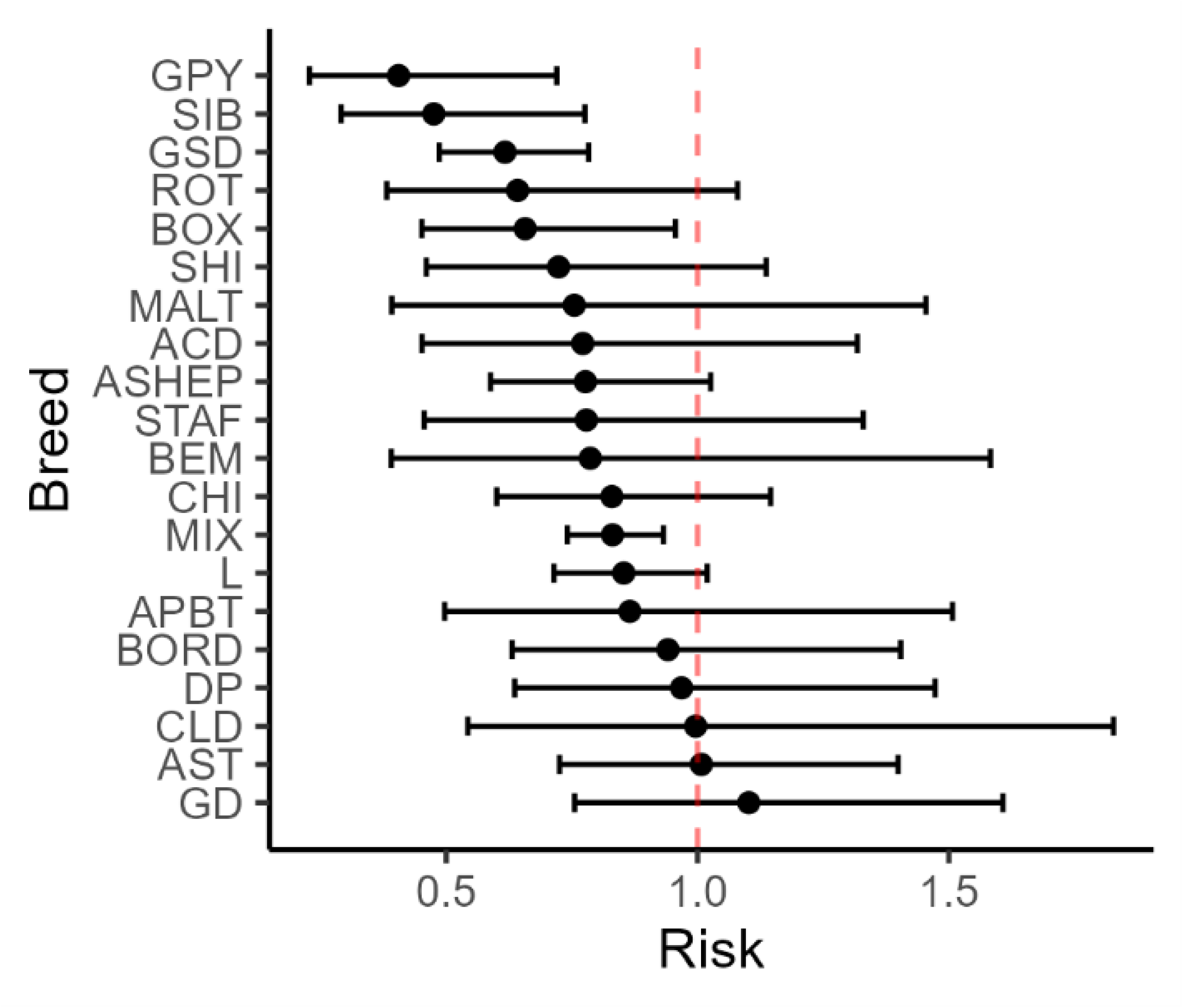
Relative risks by Bottom 20 Dog Breed for *Ancylostoma* spp.

**Figure 5:**
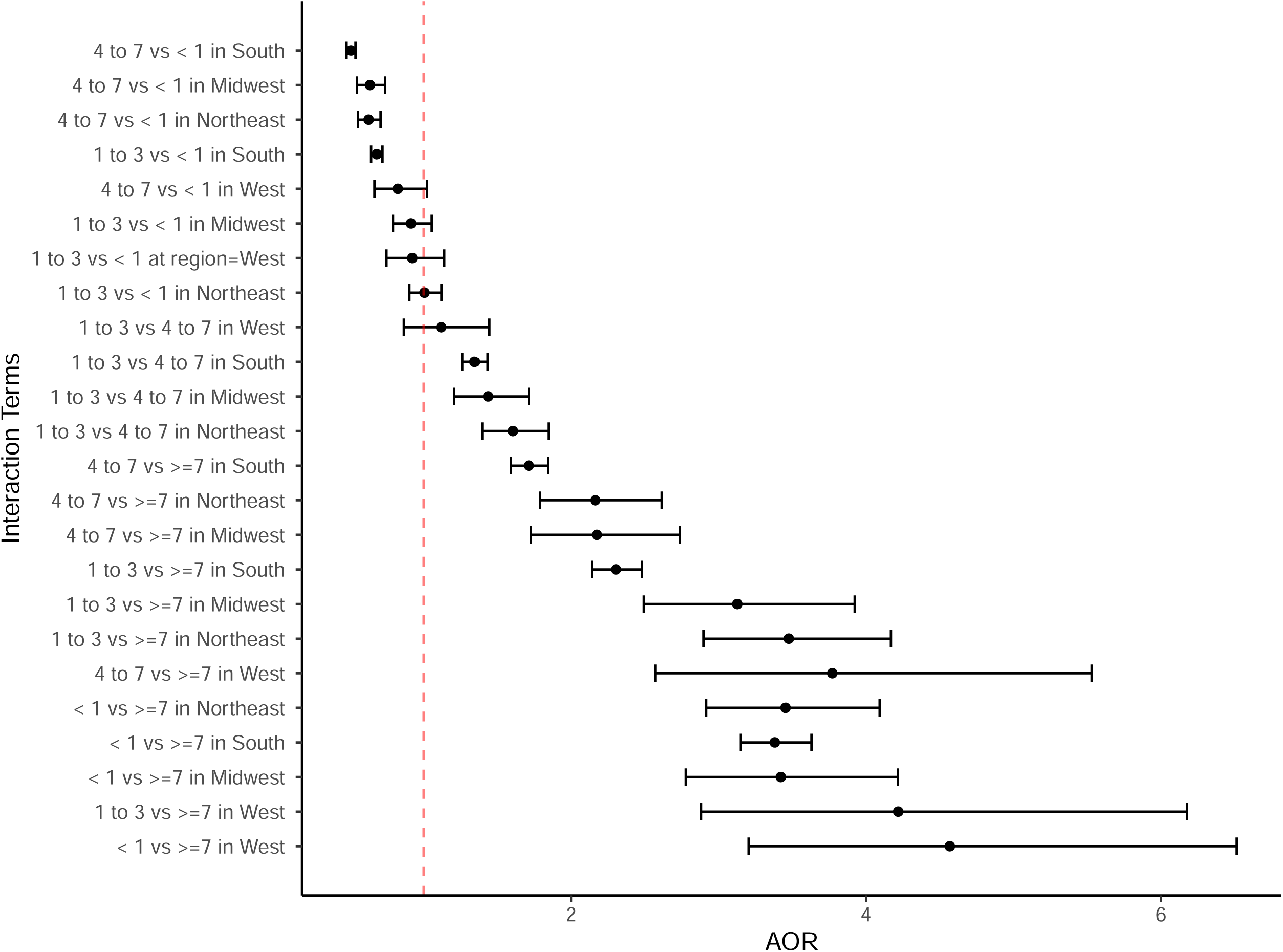
Relative risks by Bottom 20 Dog Breed for *A. caninum* F167Y

**Table 3:** The Relative Risk (RR) of *Ancylostoma* spp. and *A. caninum* F167Y for the bottom 20 breeds.

| *Ancylostoma* spp. |  |  | *A. caninum* F167Y |  |  |
| --- | --- | --- | --- | --- | --- |
| Breed | RR | 95% CI | Breed | RR | 95% CI |
| Cavalier King Charles spaniel | 0.09 | (0.06, 0.16) | Great Pyrenees | 0.40 | (0.23, 0.72) |
| Havanese | 0.15 | (0.10, 0.24) | Siberian husky | 0.48 | (0.29, 0.78) |
| Shiba inu | 0.21 | (0.11, 0.37) | German shepherd dog | 0.62 | (0.49, 0.78) |
| Bichon Frise | 0.23 | (0.15, 0.35) | Rottweiler | 0.64 | (0.38, 1.08) |
| Soft-coated wheaten terrier | 0.25 | (0.11, 0.55) | Boxer | 0.66 | (0.45, 0.96) |
| Papillon | 0.27 | (0.13, 0.54) | Shih Tzu | 0.72 | (0.46, 1.14) |
| Portuguese water dog | 0.30 | (0.14, 0.63) | Maltese | 0.75 | (0.39, 1.45) |
| Yorkshire terrier | 0.31 | (0.26, 0.38) | Australian cattle dog | 0.77 | (0.45, 1.32) |
| Lhasa Apso | 0.32 | (0.16, 0.61) | Australian shepherd | 0.78 | (0.59, 1.03) |
| Maltese | 0.33 | (0.26, 0.41) | Staffordshire bull terrier | 0.78 | (0.46, 1.33) |
| Samoyed | 0.34 | (0.15, 0.75) | Belgian Malinois | 0.79 | (0.39, 1.58) |
| Poodle, toy | 0.38 | (0.25, 0.59) | Chihuahua | 0.83 | (0.60, 1.15) |
| Poodle, miniature, +/-multicolor | 0.39 | (0.20, 0.75) | Mixed breed | 0.83 | (0.74, 0.93) |
| American Eskimo | 0.39 | (0.18, 0.86) | Labrador retriever | 0.85 | (0.71, 1.02) |
| Shih Tzu | 0.42 | (0.36, 0.49) | American pitbull terrier | 0.87 | (0.50, 1.51) |
| Pomeranian | 0.43 | (0.34, 0.55) | Border collie | 0.94 | (0.63, 1.40) |
| Scottish terrier | 0.43 | (0.21, 0.91) | Doberman pinscher | 0.97 | (0.64, 1.47) |
| Italian greyhound | 0.44 | (0.21, 0.92) | Catahoula leopard dog | 1 | (0.54, 1.83) |
| Cairn terrier | 0.45 | (0.26, 0.78) | American Staffordshire terrier | 1.01 | (0.73, 1.40) |
| West Highland white terrier | 0.48 | (0.32, 0.72) | Great Dane | 1.1 | (0.76, 1.61) |

Overall, these results underscore the importance of considering breed-specific factors in the epidemiology and management of *Ancylostoma* spp. infections along with the *A. caninum* F167Y polymorphism, which may guide targeted interventions to mitigate the burden of this parasite in canine populations.

### 3.5 Adjusted odds ratios of Ancylostoma spp. and A. caninum F167Y infections

The adjusted odds ratios (AOR) from logistic regression analysis, examining factors influencing *Ancylostoma* spp. and *A. caninum* F167Y infections, are shown in Table 4.

**Table 4:** The Adjusted Odds Ratios (AOR) from logistic regression analysis of factors influencing *Ancylostoma* spp. and *A. caninum* F167Y infections.

|  | *Ancylostoma* spp. | *A. caninum* F167Y |
| --- | --- | --- |
| Predictor variables | AOR (95% CI) | AOR (95% CI) |
| **Neuter<sup>¶</sup>** |  |  |
| Yes | 0.53 (0.511,0.56) | 0.57 (0.51, 1.64) |
| **Sex<sup>†</sup>** |  |  |
| Female | 1.04 (1.01,1.07) | 1.02 (0.94,1.11) |
| **Region<sup>§</sup>** |  |  |
| Midwest | 1.10 (0.86, 1.41) | 0.89 (0.33, 2.41) |
| South | 3.22 (2.77, 3.87) | 3.58 (1.92, 6.70) |
| West | 0.21 (0.15, 0.31) | 0.27 (0.08,0.97) |
| **Age<sup>a</sup>** |  |  |
| 1 to 3 | 3.48 (2.89, 4.17) | 7.77 (4.13,14.65) |
| 4 to 7 | 2.16 (1.79, 2.61) | 4.17 (2.16,8.03) |
| <1 | 3.45 (2.92, 4.09) | 8.10 (4.39,14.93) |

<sup>¶</sup>Reference category is No; <sup>†</sup>Reference category is Male; <sup>§</sup>Reference category is Northeast; <sup>a</sup>Reference category is $\geq 7$

#### For *Ancylostoma* spp. infections

Neutered individuals had significantly lower odds of infection compared to non-neutered individuals (AOR = 0.53, 95% CI: 0.511-0.56). Female individuals had slightly higher odds of infection compared to males (AOR = 1.04, 95% CI: 1.01-1.07). Regionally, individuals residing in the South had substantially higher odds of infection (AOR = 3.22, 95% CI: 2.77-3.87) compared to those in the Northeast, while those in the West had much lower odds (AOR = 0.21, 95% CI: 0.15-0.31).

#### For *A. caninum* F167Y infections

There were no significant differences in infection odds between neutered and non-neutered individuals (AOR = 0.57, 95% CI: 0.51-1.64). Female individuals had slightly lower odds of infection compared to males, but the difference was not significant (AOR = 1.02, 95% CI: 0.94-1.11). Regionally, individuals in the South had higher odds of infection (AOR = 3.58, 95% CI: 1.92-6.70) compared to those in the Northeast, while those in the West had lower odds (AOR = 0.27, 95% CI: 0.08-0.97).

#### Regarding age groups

Dogs aged <1 year had substantially higher odds of infection for both *Ancylostoma* spp. (AOR = 3.45, 95% CI: 2.92-4.09) and *A. caninum* F167Y (AOR = 8.10, 95% CI: 4.39-14.93) infections compared to those aged 7 years and above. Dogs aged 1 to 3 years also had significantly higher odds of infection for both *Ancylostoma* spp. (AOR = 3.48, 95% CI: 2.89-4.17) and *A. caninum* F167Y (AOR = 7.77, 95% CI: 4.13-14.65). Dogs aged 4 to 7 years had elevated odds of infection for both *Ancylostoma* spp. (AOR = 2.16, 95% CI: 1.79-2.61) and *A. caninum* F167Y (AOR = 4.17, 95% CI: 2.16-8.03) compared to those aged 7 years and above.

#### Regarding the interaction between age and region

Figure 6 represents the logistic regression analysis of the interaction between region and age influencing *Ancylostoma* spp. and *A. caninum* F167Y polymorphism infections. For *Ancylostoma* spp. infections , younger dogs (<1 year) consistently exhibited higher odds of infection across all regions when compared to older dogs (≥7 years). In the West, the odds of infection were notably higher for dogs aged <1 year compared to those aged ≥7 years (AOR = 4.57, 95% CI: 3.20, 6.51), followed closely by dogs aged 1 to 3 years (AOR = 4.22, 95% CI: 2.88, 6.18). In the Midwest, dogs aged <1 year also had significantly higher odds compared to those aged ≥7 years (AOR = 3.42, 95% CI: 2.78, 4.22), with similar trends in the South (AOR = 3.38, 95% CI: 3.15, 3.63) and Northeast (AOR = 3.45, 95% CI: 2.92, 4.09). The odds were also elevated for dogs aged 4 to 7 years compared to those aged ≥7 years across the regions, particularly in the West (AOR = 3.77, 95% CI: 2.57, 5.53).

**Figure 6:**
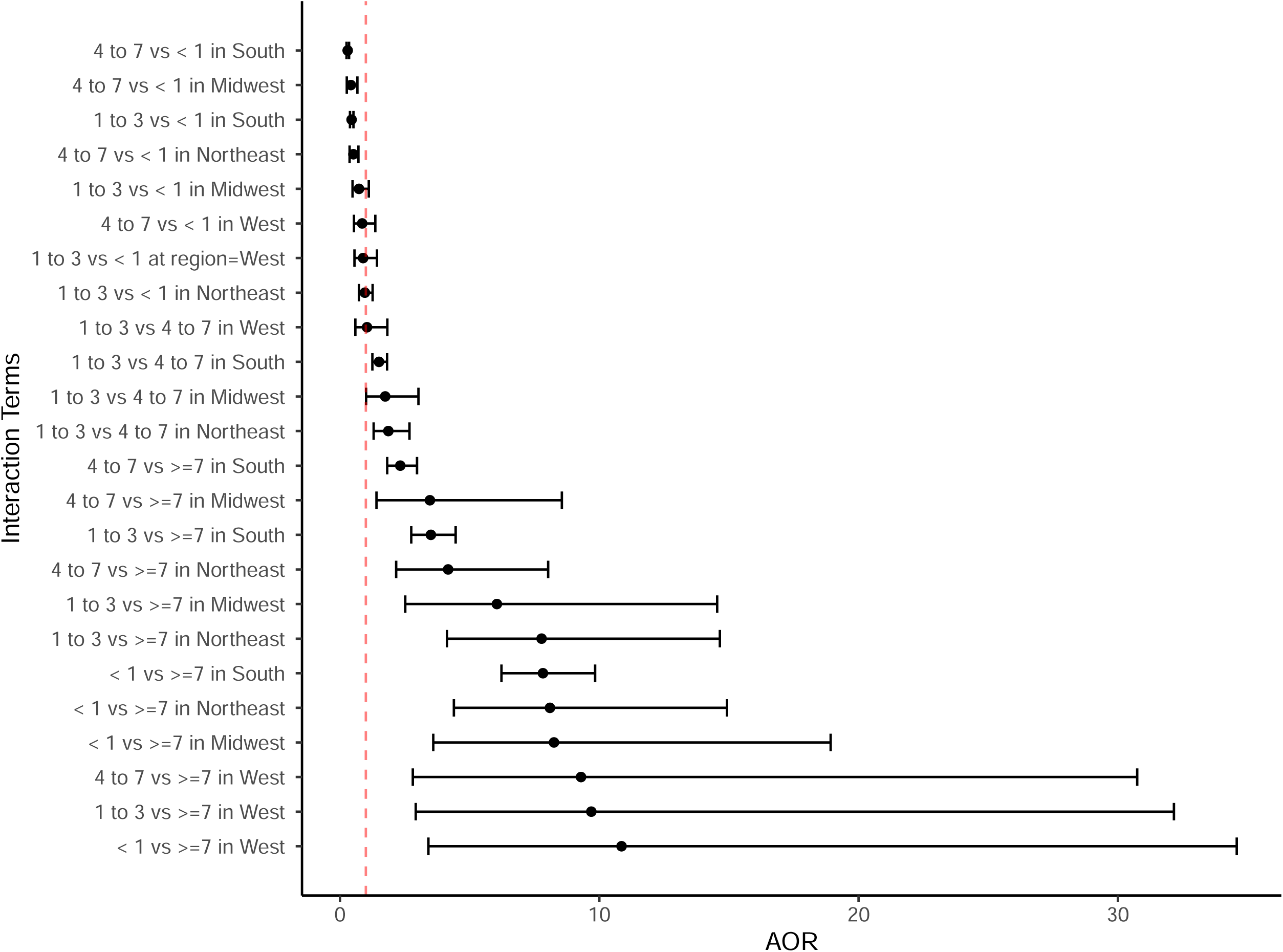
A. Adjusted Odds Ratios (AOR) for interaction terms between region and age from logistic regression analysis of factors influencing *Ancylostoma* spp. infections. B: Adjusted Odds Ratios (AOR) for interaction terms between region and age from logistic regression analysis of factors influencing *A. caninum* F167Y infections.

For infections with the *A. caninum* F167Y, younger dogs generally had higher odds of infection, especially in the West, where dogs aged <1 year had the highest odds compared to those aged ≥7 years (AOR = 10.86, 95% CI: 3.41, 34.59), and dogs aged 1 to 3 years also showed elevated odds (AOR = 9.69, 95% CI: 2.92, 32.16). Similar trends were observed in the Midwest, where dogs aged <1 year exhibited significantly higher odds compared to those aged ≥7 years (AOR = 8.25, 95% CI: 3.60, 18.93), and in the Northeast (AOR = 8.10, 95% CI: 4.39, 14.93). The South also demonstrated elevated odds for younger dogs, particularly for those aged <1 year compared to those aged ≥7 years (AOR = 7.83, 95% CI: 6.23, 9.84). The odds for dogs aged 4 to 7 years compared to those aged <1 year were lower across regions, with the lowest in the Midwest (AOR = 0.42, 95% CI: 0.26, 0.67) and the South (AOR = 0.30, 95% CI: 0.25, 0.35).

Overall, these findings suggest that neuter, sex, region of residence, and age are significant factors influencing the prevalence of *Ancylostoma* spp. and *A. caninum* F167Y infections but are not equivalent between susceptible and resistant populations.

## 4. Discussion

To the authors’ knowledge, this is the first study evaluating risk association and odds ratio analyses between patient characteristics (breed, neuter status, age, geographical location, and sex) and detection of *Ancylostoma* spp. and *A. caninum* F167Y polymorphism in dogs using a commercially available molecular qPCR test. Data from a total of 885,424 canine fecal samples collected between March 2022 and December 2023 from the US were included in the analysis, of which 15,537 were detected for *Ancylostoma* spp. (1.90%). In this canine hookworm population, the F167Y polymorphism associated with benzimidazole resistance was detected in 2,244 samples (14.44%). Of these, a subset of positive canine results for *Ancylostoma* spp. (n=15,190) and *A. caninum* F167Y (n=2,244) were selected for the association analysis.

Puppies (aged less than 1 year) exhibited the highest prevalence and significantly elevated AOR for both *Ancylostoma* spp. detection and the *A. caninum* F167Y polymorphism, 3.71% and 4.46, and 17.13% and 2.59, respectively. This observation aligns with a recent study where, out of 206 dogs with age-related information, the prevalence of the F167Y polymorphism was notably higher in puppies compared to seniors (Venkatesan et al., 2023). Additionally, our previous research findings also support this trend, revealing a prevalence of 3.8% and 14.5% for *Ancylostoma* spp. and *A. caninum* F167Y, respectively (Leutenegger et al., 2024). The increased use of anthelmintic treatments recommended for lactating dams until whelping and puppies during their first year can contribute to the selection pressure favoring resistant survivors (Von Samson-Himmelstjerna et al., 2021). The intervals between these treatments are often shorter than the pre-patent period for hookworms, minimizing biodiversity and the availability of refugia in the treated individual. Consequently, genetically resistant worms that survive treatment gain a significant reproductive advantage, and the scarcity of refugia leads to a rapid increase in their frequency (Martin et al., 1981; van Wyk, 2001). These combined factors exert substantial selection pressure for drug resistance in nematodes, mirroring the epidemiological conditions that have driven high levels of MADR in gastrointestinal nematodes of sheep and goats worldwide (Wolstenholme et al., 2004; Kaplan and Vidyashankar, 2012).

Furthermore, upon incorporating the data from 2023, we observed a shift in the submission population. Specifically, there is an increased number of samples from dogs categorized as ‘mature adults’ and a decreased number of samples from puppies and may represent a shift in usage of the parasite panel from a more disease-related to wellness-related tool. This comparison is made against the total submissions reported in our previous publication (Leutenegger et al., 2024). Consequentially, the data will change over time with shifting submission demographics. This aspect needs to be considered in the interpretation of the data with the continued collection of parasite screening results.

Interestingly, the West region had the highest prevalence for the *A. caninum* F167Y of 19.78%, across the regions. This is in line with findings from previous studies using a different molecular approach with next generation sequencing, where the prevalence (75%) and mean allele frequency of the F167Y SNP (72.8%) for this region was significantly higher compared to other regions of the US (Venkatesan et al., 2023). This corroborates our previous findings as well, where the West showed the highest prevalence for *A. caninum* F167Y polymorphism (13.4%) (Leutenegger et al., 2024, Leutenegger et al., 2023a). One possible explanation for the higher resistance prevalence is that there is a lower level of environmental refugia in the west due to the drier climate. Low levels of refugia lead to higher drug selection pressure and is thus recognized as an extremely important factor for the development of anthelmintic resistance in nematodes of livestock. This circumstance would therefore parallel the situation with *H*. *contortus* in sheep in drier and/or colder climates (Queiroz et al., 2020; Papadopoulos et al., 2001)

Based on the previous body of literature, it was expected for greyhounds to have the highest RR for hookworm infection (8.03). A recent investigation examined the likelihood of positive results for hookworms in greyhounds compared to other breeds. The study analyzed fecal samples using two methods: fecal flotation with zinc centrifugation and fecal coproantigen testing. The odds of detecting hookworms were 15.3 using the flotation method and 14.3 using the antigen test in greyhounds (Burton et al., 2024). Greyhounds exhibit several breed-related differences. Research has shown that greyhounds have lower leukocyte (Porter and Canaday, 1971) and neutrophil counts (Steiss et al., 2000) compared to the reference values for canines. Additionally, greyhounds are hypoproteinemic relative to other dog breeds due to low serum α and β-globulin concentrations (Fayos et al., 2005). These factors may potentially influence the susceptibility of greyhounds to nematode infection when compared to other breeds. However, further studies are needed to confirm this hypothesis.

Interestingly, greyhounds were not the breed at the highest risk for *A. caninum* F167Y SNP detection [fourth highest RR (2.45)] and rather old English sheepdogs, American foxhounds, and toy poodles, were the top three breeds at risk for the detection of hookworms with the resistant polymorphism. This is in line with our previous work where poodles, Bernese Mountain dogs, and cocker spaniels had a higher prevalence than greyhounds for the F167Y polymorphism associated with benzimidazole resistance (Leutenegger et al., 2024). The finding becomes even more intriguing when considering recent research. In a study involving 35 foxhounds from a New Jersey kennel with a history of persistent *A. caninum* infections, despite anthelmintic treatment, the efficacy of three commercial anthelmintic products was evaluated using fecal egg count reduction (FECR) at 11 days post-treatment. The authors observed a reduced efficacy of <70% across the major drug classes (benzimidazoles, macrocyclic lactones, and pyrantel) (Balk et al., 2023). Similarly, another study conducted in a Labrador retriever kennel in Georgia showed reductions in FECR at 11 days post-treatment (<60%) for these same drug classes (Jimenez Castro et al., 2022). Based on these FECR data, the most likely cause of the persistent egg shedding was MADR in both studies (Jimenez Castro et al., 2020). Interestingly, there was no known interaction with greyhounds, and no greyhounds had ever been on the premises of either kennel. Taking this into account, it is possible that the source of these MADR infections originated from within each kennel and subsequently spread due to ongoing selection pressure from treatment. However, DNA sequencing of the β-tubulin gene from pre- and post-treatment hookworm eggs in the fenbendazole treatment group of the Labrador retriever kennel study from Georgia revealed that the F167Y and Q134H mutations were present on the same β-tubulin haplotypes found in MADR *A. caninum* recovered from 154 greyhounds residing in kennels across 16 different locations in 8 US states (Jimenez Castro et al., 2021). These genetic data suggest that the MADR worms in the Labrador retriever kennel were almost certainly introduced by a dog infected elsewhere with worms that originally derived from a greyhound (Jimenez Castro et al., 2022). Molecular studies were not performed in the Foxhound kennel study to test this hypothesis. Further studies are needed to explore the molecular epidemiology of MADR *A. caninum* infections in non-greyhound kennels to evaluate the non-greyhound origin of MADR.

Intact dogs had a higher prevalence of 3.88% and 15.66% compared to neutered dogs, for hookworm infection and *A. caninum* F167Y, respectively. This could be due to the fact that the majority of neutered dogs are medicalized and receive heartworm preventives which most of these have a hookworm efficacy claim. However, few studies have specifically explored the link between neuter status and hookworm infection. For instance, a recent study analyzed fecal samples from 3,022 dogs across 288 dog parks in the US. The study found that 89.8% of female dogs were spayed, and 84.6% of male dogs were neutered. Unfortunately, the authors did not provide parasite prevalence data based on these patient characteristics (Stafford et al., 2020). Similarly, another study conducted in France examined samples from 414 pet dogs during 2017-2018. The results showed a higher overall parasite prevalence in intact dogs (14.8%) compared to neutered dogs (7.5%). However, the authors did not specify the parasite species responsible for this difference (Bourgoin et al., 2022). Additionally, reproductive status plays a role in a dog’s behavior, affecting their willingness to obey commands. This is particularly relevant for working dogs, such as those in hunting kennels. These dogs may be subject to an increased risk of exposure to hookworm infective larvae compared to neutered dogs, whether in wildlife reservoirs or contaminated kennel environments (Serpell et al., 2005). Interestingly, in a study looking into the effects of sterilization not only on longevity, but also on pathophysiological causes of death evaluating health records from 41,045 dogs between 1984 and 2004, the authors found that lifespan was greater in the neutered dogs compared to intact subjects. Furthermore, of five infectious diseases that were considered, four had significantly lower frequencies in sterilized dogs, one of these being intestinal parasites albeit no specific etiological agent (Hoffman et al., 2013).

Overall, these findings provide valuable insights into the demographic and regional factors influencing the prevalence of *Ancylostoma* spp. infections and the distribution of the *A. caninum* F167Y polymorphism in dogs, which may inform targeted prevention and control strategies for veterinarians.

This study had limitations common for retrospective studies and lack of corresponding clinical information. Important information such as home environment, clinical history, presentation, and clinical signs or if for routine wellness visit, as well as travel and anthelmintic history were unavailable for this population. Furthermore, for the neutering status analysis we were not able to account if the unneutered dogs were to become neutered in the future. Also, even though we used the ZIP codes from the place where the sample was submitted to assign this to one of the four regions, this does not necessarily mean that the infection was acquired in that geographical region. Furthermore, the F167Y polymorphism could have been detected in other hookworm species infecting dogs in the US, such as *Ancylostoma braziliense* or *Uncinaria stenocephala*, even though recent data has shown that none of the benzimidazole resistance-associated polymorphisms were detected (Stocker et al., 2024) in the same sample set analyzed by Venkatesan et al., 2023. In addition, the recently identified Q134H SNP linked to benzimidazole resistance (reported by Venkatesan et al., 2023) and subsequently incorporated into the qPCR test during the summer of 2023, was not part of the dataset analyzed in this study. This finding highlights the ongoing evolution of drug resistance mechanisms in *A. caninum* and underscores the importance of monitoring genetic changes in this parasite.

## 5. Conclusions

Our study highlights the utility of an accessible diagnostic qPCR parasite panel with fast turnaround times to provide insights into the molecular epidemiology of hookworm and benzimidazole resistance associated with the F167Y polymorphism. This research sheds light on the fascinating interplay between genetics, treatment efficacy, and the spread of drug-resistant hookworms in canine populations. This study establishes a foundation for continuous surveillance initiatives, promotes awareness of One Health and antimicrobial stewardship, and emphasizes risk factors and odds ratios that inform clinical decision-making and enhance the efficient utilization of anthelmintic drugs in companion animals.

## Acknowledgements

We would like to thank the parasitology technicians for performing the parasitology protocols. This research did not receive any specific grant from funding agencies in the public, commercial, or not-for-profit sectors. All research, work, materials, and medical writing was funded through Antech Diagnostics internal mechanisms. The commercial real-time qPCR test used for analysis of samples in this study was KeyScreen^®^ GI Parasite PCR, an Antech Diagnostics (Mars Petcare Science & Diagnostics) product.

## Authors’ contributions

PDJC, CML, CEL, JLW, were involved in study inception, PDJC, CML, HR, HLR, HEM, and JLW developed the study design, data analyses, interpretation, and statistics; CML directed the study. All authors contributed to the writing and editing of the manuscript. All authors have read and approved the final manuscript.

